# Aging remodels the human muscle secretome, impairing stem cell via sphingolipid metabolism

**DOI:** 10.64898/2026.10.01.755640

**Authors:** Barbara Crisol, Natalia Pinzón, Mona Bensalah, Jessica Ohana, Marie Lhomme, Maharajh Ponnaiah, Johanna Bruce, Maria Kondili, Ludovic Gaut, Elisa Negroni, Alexis Boulinguiez, Gillian Butler-Browne, Vincent Mouly, Capucine Trollet, Anne Bigot

**Author notes:** Co-last and co-corresponding authors and.

## Abstract

Aging reduces skeletal muscle regenerative capacity, but how the aged environment impairs muscle stem cells (MuSCs) remains unclear. Skeletal muscle acts as a secretory organ, yet age-related changes in its secretome and impact on stem cells are poorly understood.

We performed proteomic and lipidomic analyses of conditioned media from primary muscle cells of young (15–25 years) and aged (60–86 years) donors. We identified 127 age-dependent secreted proteins, revealing major remodeling. The lysosomal enzyme acid sphingomyelinase (ASM), which converts sphingomyelin into ceramide, was the most upregulated factor.

Recombinant ASM reduced PAX7+ MuSCs in young cultures, showing extracellular ASM impairs stem cell maintenance. Lipidomics revealed disruption of the sphingomyelin–ceramide–S1P axis, with decreased sphingomyelin and S1P, a lipid promoting MuSC proliferation. Exogenous S1P restored the stem cell pool in aged cultures.

These findings link secretome remodeling and sphingolipid metabolism to MuSC decline and identify ASM and S1P as targets.

## Introduction

Skeletal muscle plays a crucial role in body locomotion, posture and metabolism, accounting for approximately 40% of total body mass and nearly 30% of whole-body metabolic rate in adult humans (Frontera & Ochala 2015). Skeletal muscle is composed of multinucleated contractile myofibers surrounded by a heterogeneous population of mononuclear cells, including muscle stem cells (satellite cells, MuSCs), fibroadipogenic progenitors (FAPs), endothelial cells, immune cells and motoneurons. Beyond its contractile function, skeletal muscle acts as a secretory organ, releasing a wide range of bioactive factors that regulate muscle homeostasis and mediate communication with local and distant cell populations (Severinsen & Pedersen 2020). This coordinated interplay between muscle fibers and their surrounding cell populations is essential for maintaining tissue integrity and ensuring efficient regeneration following injury (Fuchs & Blau 2020; Rodríguez et al. 2024).

With age, skeletal muscle regenerative capacity following injury declines and is often accompanied by a progressive loss of muscle mass (Pollina & Brunet 2011). When this loss of mass, strength, and function reaches a pathological threshold, it is defined as sarcopenia, a condition that significantly impacts health and survival in older individuals (Srikanthan & Karlamangla 2014). Aging is also associated with a chronic low-grade inflammatory state, termed inflammaging, characterized by the accumulation of pro-inflammatory factors and cellular damage. While the decline in muscle regenerative capacity has often been attributed to alterations in MuSCs (Sousa-Victor et al. 2014), both intrinsic and extrinsic mechanisms have been implicated (Relaix et al. 2021). In mice, aging has been associated with increased MuSC senescence and functional decline (Sousa-Victor et al. 2014), supporting a role for cell-intrinsic alterations, and, recent work has shown that senescent cells contribute to the establishment of an inflamed niche that impairs muscle regeneration (Moiseeva et al. 2023). Early studies using heterochronic parabiosis and transplantation approaches have also demonstrated that exposure to a young systemic or local environment can restore regenerative capacity in aged MuSC (Carlson & Faulkner 1989; Conboy et al. 2005) highlighting the influence of extrinsic signals (Sousa-Victor et al. 2022). Consistently, studies in human systems indicate that aged MuSCs can retain substantial regenerative potential (Bigot et al. 2015; Novak et al. 2021), further supporting the idea that age-related impairments are not solely cell-autonomous. Together, these finding underscore the importance of the tissue environment and intercellular communication in regulating MuSC function during aging (Fuchs & Blau 2020; Hong et al. 2022; Rodríguez et al. 2024).

In this context, muscle-derived secreted factors (Severinsen & Pedersen 2020) have emerged as important regulators of tissue homeostasis. While numerous protein mediators, including myokines, have been identified, how aging remodels the muscle secretome and whether these changes directly regulate stem cell function remain poorly understood. In particular, the contribution of lipid-mediated intercellular signaling remains poorly defined. Muscle cells release not only soluble factors but also extracellular vesicles (EVs), which carry proteins, nucleic acids, metabolites and lipids to recipient cells (Le Bihan et al. 2012). Among these components, lipids are increasingly recognized as key regulators of EV function and intercellular communication (Rome & Tacconi 2024).

Sphingolipids constitute a major class of bioactive lipids involved in the regulation of cell fate, inflammation, and stem cell activity (Hannun & Obeid 2018). Within this network, Sphingosine-1-phosphate (S1P) has been shown to promote MuSC activation and muscle regeneration (Calise et al. 2012; Cordeiro et al. 2019; Hakkar et al. 2025; Nagata et al. 2006). S1P production is tightly linked to sphingomyelin metabolism, in which sphingomyelin hydrolysis generates ceramide, a bioactive lipid that can be further converted into sphingosine and S1P. Acid sphingomyelinase (ASM), a key enzyme controlling sphingomyelin turnover, has been implicated in aging and inflammatory processes across multiple tissues (Park et al. 2018; Tam et al. 2010). However, whether and how these pathways are altered in aging skeletal muscle, particularly at the level of the muscle secretome, and how they influence MuSC function, remains unknown.

Here, we investigated how aging remodels the human skeletal muscle secretome, with a particular focus on its lipid components. By combining proteomic and lipidomic analyses of primary muscle cells from young and aged donors, and integrating these data with human plasma aging signatures, we identify alterations in lysosome-related pathways, including dysregulation of the sphingomyelin-ceramide-S1P axis and increased secretion of ASM as features of aged muscle cells. Functionally, these changes impair MuSC maintenance and regenerative potential, revealing a role for lipid-mediated intercellular communication in linking secretome remodeling to stem cell dysfunction during human muscle aging.

## Results

### Myotubes from aged donors retain hallmarks of aging

To investigate age-related alterations in myogenic potential and secretory profile of human skeletal muscle, we used an *in vitro* model based on the isolation and differentiation of skeletal muscle stem cells (MuSC) from young (15-24 years) and aged (60-86 years) donors (Fig. 1a), with “young” and “aged” referring throughout the manuscript to cells derived from these respective donor groups. While this *in vitro* system does not recapitulate the full complexity of injury-induced regeneration, it allows assessment of the early stages of regeneration in terms of cell-autonomous myogenic differentiation and secretion. To ensure homogeneous myogenic purity in each culture, the percentage of myoblasts (Desmin-positive cells) was monitored in all cultures. When necessary, cultures were enriched by MACS sorting using CD56 to increase the proportion of Desmin-positive cells (Bensalah et al. 2022). All cultures contained at least 82% Desmin positive cells (Fig S1a). Consistent with our previous observations (Bigot et al. 2015), donor age did not affect cell proliferation (Fig. 1b and S1b) or myoblast fusion capacity (Fig. 1c). After three days of differentiation, cultures derived from aged donors contained fewer PAX7+ cells, a marker of quiescent MuSC, compared with cultures derived from young donors (Fig. 1d).

**Fig 1:**
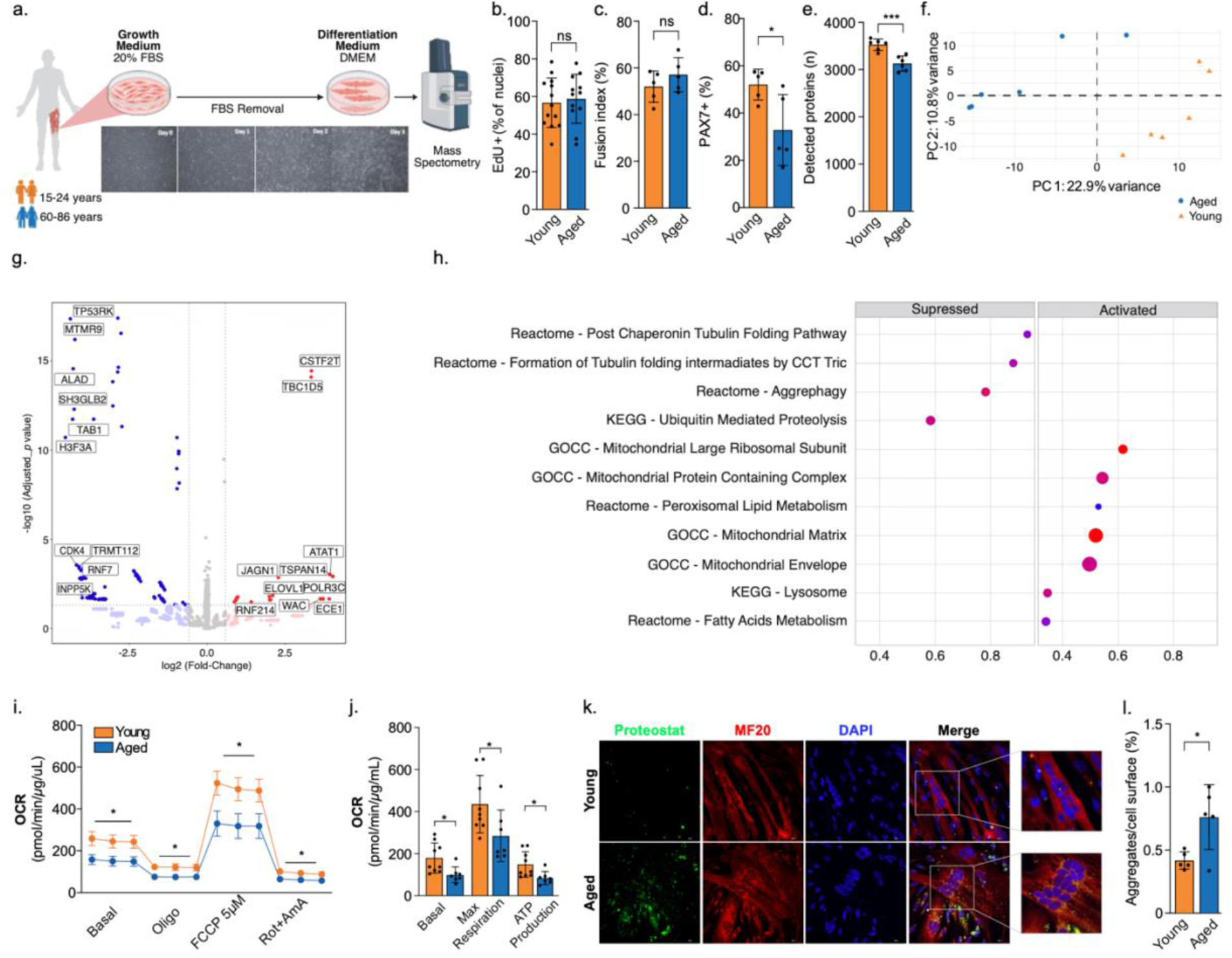
Muscle cells from aged donors retain aging-associated hallmarks. **a**,Differentiation protocol of skeletal muscle cells. Myotubes were collected and analyzed by mass spectrometry. **b,** Analysis of Edu+ myoblasts from young and aged cultures, n=14-15 cell lines. **c-l** Analyses were performed in primary myotubes differentiated for 3 days from young and aged donors. **c,** Fusion index analysis, n=5 donors. Data are presented as mean ± s.d. **d,** PAX7 positive (PAX7+) cells in the cultures of young and aged donors, n=5. **e,** Number of different detected proteins on the proteome analysis, n=6. Data are presented as mean ± s.d. **f,** PCA of quantitative proteome data from young and aged myotubes. Each dot corresponds to one sample. Young (orange triangles) and aged (blue dots) groups show separation along the principal components, reflecting age-dependent remodeling of the muscle cell proteome, n=6. **g,** Volcano plot of differential protein expression between young and aged myotubes. Each point represents a quantified protein, plotted by log₂ fold change (aged vs. young) and −log₁₀ adjusted p-value. Proteins significantly up (red) or downregulated (blue) with age are highlighted. The Top 20 proteins modulated by age are named in the graph. **h,** Enrichment analysis of proteomic data comparing young and aged myotubes. Selected enriched pathways are ranked according to normalized enrichment score (NES), with significance assessed by false discovery rate (FDR). Pathways positively enriched (activated) and reduced (suppressed) in aged myotubes are indicated. **i,** Oxygen consumption rate (OCR) during a mitochondrial stress test, n=7-9. Data are presented as mean +SEM. **j,** OCR from basal, maximum and APT-linked respiration during mitochondria stress test n=7-9. Data are presented as mean ± s.d. **k,** Representative confocal images from primary human myotubes from young and aged donors (scale bar, 50 µm) stained with the Proteostat fluorescent dye (green) and MF20 (red). **l,** Aggregates per cell surface in young and aged myotubes, n=5. Data are presented as mean ± s.d. ns not significant; *p<0.05 vs. young; ***p<0.001 vs young.

To further validate that myotubes differentiated from aged donors retain aging features *in vitro*, we performed proteomic analyses of three-day differentiated myotubes (Fig. 1a and S1c. Table S1). Although equal amounts of total protein were analyzed for each sample, the proteome of myotubes from aged donors showed a lower number of different detected proteins per sample compared to that of myotubes from young donors (Fig. 1e), indicating reduced proteomic diversity in the proteins expressed by aged myotubes. Unsupervised principal component analysis (PCA) of proteomic profiles showed a clear separation between myotubes from young and aged donors (Fig. 1f). In total, 408 proteins were differentially expressed of which 222 were downregulated and 186 were upregulated with age (Fig. 1g and S1d). Gene set enrichment analysis (GSEA) performed on the proteomic dataset revealed in aged myotubes down-regulation of proteostasis-related pathways, including aggrephagy and ubiquitin-mediated proteolysis, and enrichment in mitochondrial complex proteins (Fig. 1h). Enrichment analysis also highlighted an activation of peroxisomal lipid metabolism, a pathway involved in long-chain fatty acid degradation and phospholipid synthesis (Fig. 1h). Overall, these alterations confirm in our *in vitro* model well-established hallmarks of muscle aging, including impaired proteostasis and metabolic remodeling, consistent with observations in aged human skeletal muscle *in vivo* (Lima et al. 2023). To further confirm these predicted aging features, we next assessed mitochondrial function and metabolic parameters in differentiated myotubes. Aged myotubes exhibited impaired mitochondrial metabolism (Fig. 1i), including reduced basal respiration capacity, reduced maximal respiratory capacity and ATP production (Fig. 1j), as previously described *in vivo* (Gonzalez-Freire et al. 2018). In addition, aged myotubes displayed a lower proton leak compared to young myotubes (Fig. S1e), consistent with a reduced basal metabolic rate (BMR), a known feature of aging (Pontzer et al. 2021). These data suggest a compensatory response to the reduced mitochondrial activity by increasing level of mitochondrial proteins as observed in the enrichment analysis (Fig. 1h). Aging is also characterized by a loss of proteostasis, leading to the accumulation of misfolded proteins and increased protein aggregation (Vilchez et al. 2014). We observed that myotubes from the aged donors contained significantly higher content of protein aggregates compared to those from young donors (Fig. 1k-l). Together, these data demonstrate that muscle progenitor cells differentiated *in vitro* retain intrinsic aging-associated characteristics.

### Remodeling of the aged muscle secretome identifies ASM as a key sphingolipid-related secreted factor

To address how aging can affect muscle secretion, we performed proteomic analysis of the conditioned media - called secretome - from young and aged myotubes (Fig. 2a). Proteomic analysis of these conditioned media revealed no significant difference in the total number of different secreted proteins between young and aged myotubes (Fig. 2b and Table S2). However, 127 proteins were differentially secreted (Fig. 2c and S1d), with 102 upregulated and 25 downregulated proteins in the aged group. PCA demonstrated an age-dependent separation of the muscle secretome (Fig. S2a). Notably, a larger proportion of proteins were over-secreted in aged myotubes (Fig. 2c and Fig. S1d). Over-representation analysis (ORA) using STRING and REACTOME annotations revealed significant enrichment of sphingolipid and glycosphingolipid metabolism pathways in the aged secretome (Fig. 2d). Among the over-secreted proteins, acid sphingomyelinase (ASM), a lysosomal enzyme that converts sphingomyelin into ceramides, was the most strongly increased in aged secretome (Fig. 2c). Notably, ceramide accumulation has been previously reported during aging and is known to impair muscle metabolism (Lima et al. 2023; Laurila et al. 2022). To determine whether increased ASM secretion affects MuSC behavior, we treated 2 days differentiated myotubes derived from young donors with recombinant human ASM (Fig. 2e). ASM treatment significantly reduced the number of PAX7+ reserve cells in these young cultures (Fig. 2f-g), without affecting the fusion index (Fig. 2h), demonstrating that extracellular ASM negatively regulates MuSCs pool size, thereby recapitulating an aging-like phenotype. Together, these results identify ASM as a functional mediator of age-associated secretome remodeling.

**Fig 2:**
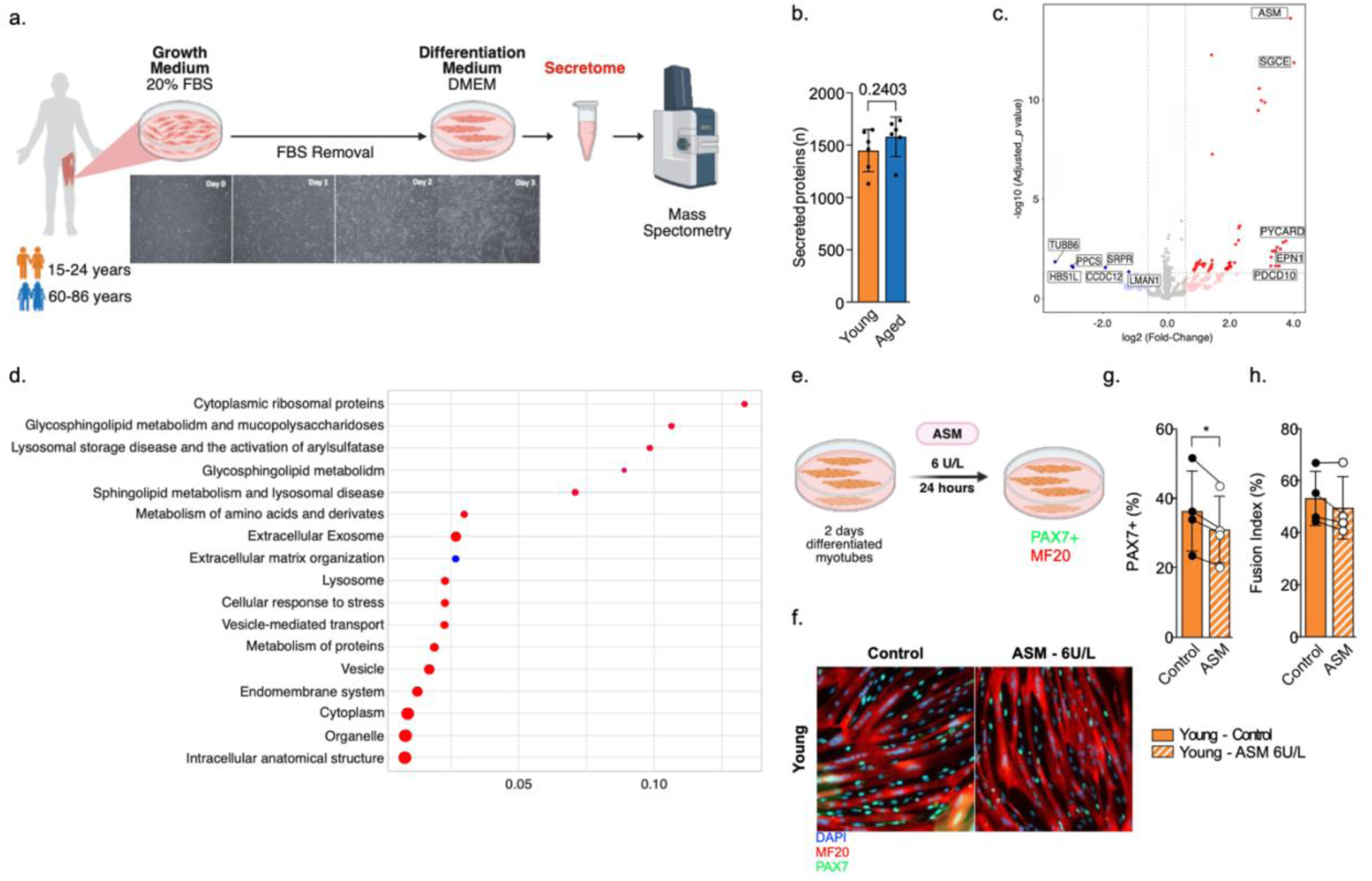
Aging effects in the secretome from primary human myotubes. **a**,Scheme of the protocol for the secretome collection. Primary cells were isolated from muscle biopsies from young and aged donors. Myoblasts were differentiated during 3 days in a serum-free media environment. Conditioned media (secretome) were collected and analyzed by mass-spectrometry **b,** Number of detected proteins on the secretome analysis, n=6. Data are presented as mean ± s.d. **c,** Volcano plot of differential proteins secreted between young and aged conditioned media. Each point represents a quantified protein, plotted by log₂ fold change (aged vs. young) and −log₁₀ adjusted p-value. Proteins significantly up (red)- or downregulated (blue) with age are highlighted. The name of the 11 mostly affected by aging are presented in the graph. **d,** Over-expression analysis (ORA) from the muscle secretome from young and aged donors. Dot size corresponds to the number of proteins associated to each pathway. Dot color indicates the adjusted p-value. Pathways are ranked based on enrichment significance. **e,** Schematic representation of the experimental setup. Myotubes differentiated for 2 days were treated with acid sphingomyelinase (ASM, 6 U/L) for 24 hours prior to analysis. **f,** Representative images from young myotubes after ASM 6U/L treatment stained for PAX7 (green), MF20 (red) and DAPI (blue). **g,** PAX7 positive (PAX7+) cells in the young culture after ASM treatment. n=4 cell lines. **h,** Fusion index analysis after ASM treatment, n=4 cell lines. Data are presented as mean ± s.d. *p<0.05.

### Aging promotes coordinated lysosomal proteins secretion required for membrane integrity

Because ASM is a lysosomal enzyme secreted via Ca^2+^-dependent lysosomal exocytosis (Tam et al. 2010), we next investigated whether aging promotes coordinated lysosomal secretion in human myotubes. Comparison of our proteins of interest (POI) from the secretome data with a reference lysosomal proteome (total 404 proteins) from human cell lines (Akter et al. 2023), revealed that, of the 127 age-regulated proteins detected in the muscle secretome, 13 were lysosomal proteins and were upregulated in the aged myotubes secretome (Fig. 3a-b). Interestingly when we compared the biological processes enriched in our muscle secretome with those identified in a systematic literature review of human aging plasma biomarkers (Moaddel et al. 2021), we observed that the lysosomal process was shared between the two datasets (Fig. 3c). Since lysosomal exocytosis is a key mechanism involved in plasma membrane repair through the extracellular release of ASM (Corrotte et al. 2013; Tam et al. 2010), we next tested whether ASM activity contributes to membrane integrity in aged myotubes. We exposed young and aged myotubes to desipramine (DPA), a functional inhibitor of lysosomal ASM, at a concentration previously shown to efficiently inhibit ASM activity without overt toxicity (30 µM) (Fig. 3d). Aged myotubes under ASM inhibition presented an increased intracellular influx of FM1-43 (Fig. 3e-f), indicating that ASM secretion is specifically required to maintain membrane integrity in aged myotubes. Importantly, without ASM inhibition aged myotubes were able to efficiently reseal their plasma membrane even following exposure to the pore-forming toxin streptolysin-O (SLO), as shown by the exclusion of FM1-43 dye (Fig S3a-b). These findings suggest that aged myotubes are not intrinsically defective in membrane repair, but rather rely on increased ASM secretion as a compensatory mechanism to preserve membrane integrity during aging.

**Fig 3:**
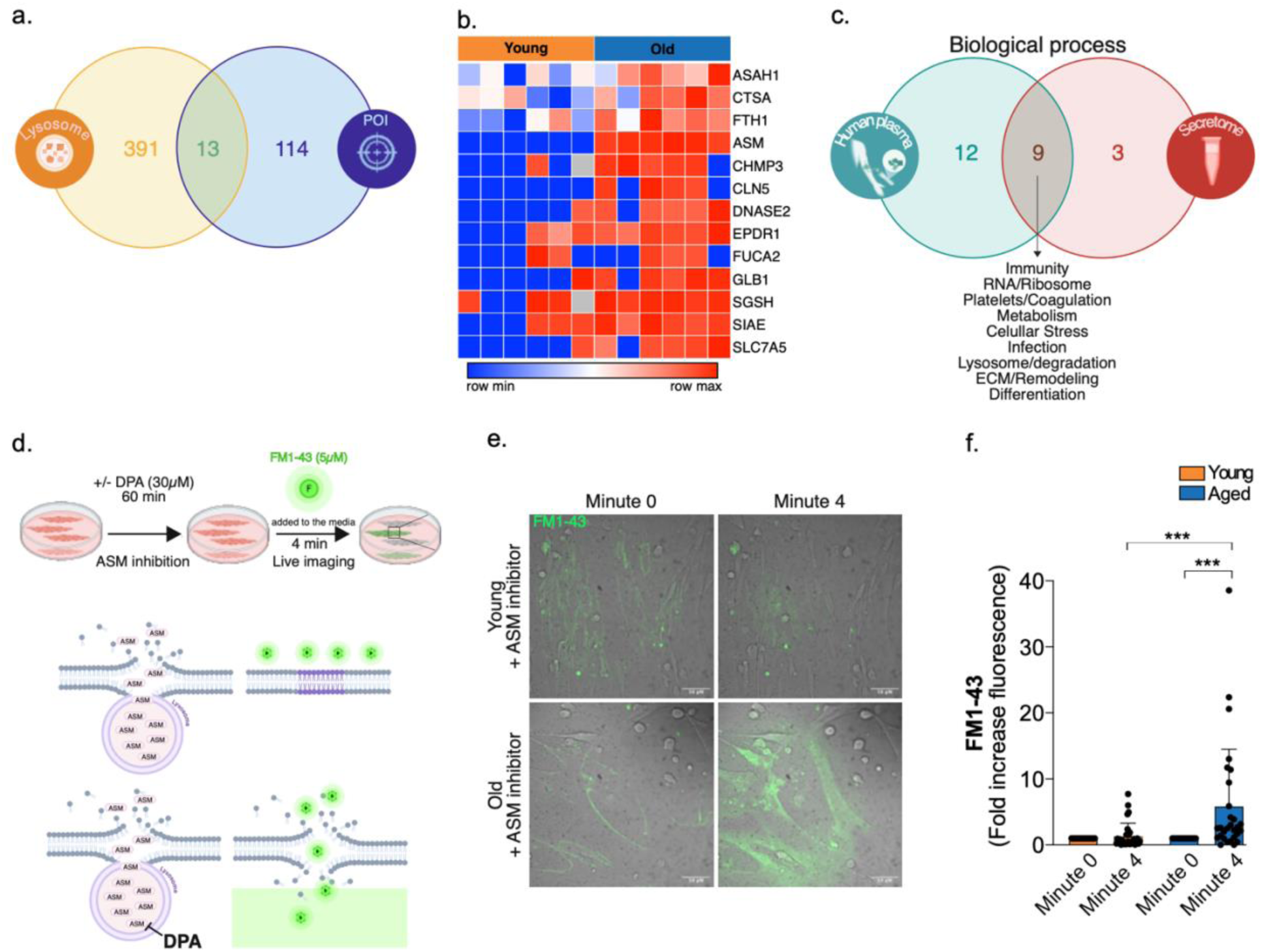
Aging increased lysosomal secretion to preserve membrane integrity. **a**,Venn diagram illustrating the overlap among the proteins of interest from the secretome (POI – blue) and the lysosomal proteomic (lysosome - yellow). The numbers indicate the counts of proteins unique to each dataset as well as those shared **b,** Heatmap of the 13 lysosomal POI upregulated in the human muscle secretome. Rows are the proteins and columns are the samples. Color scale represents z- scored abundance (blue = low, red=high), n=6. **c,** Venn diagram showing the overlap between the biological pathways from aging-associated proteins reported in the human plasma and biological pathways from proteins secreted by aged myotubes. Nine pathways were shared between both lists**. d,** Schematic representation of the experimental setup. Myotubes differentiated for 3 days were exposed to DPA (30µM) for 60 minutes prior to analysis. Subsequently, FM1-43 dye was added to the media, and the myotubes were imaged during 4 minutes. Under conditions of membrane damage accompanied by ASM secretion, membrane repair occurs efficiently, preventing FM1-43 entry into the intracellular space. In contrast, upon ASM inhibition by DPA, membrane repair is impaired, allowing FM1-43 dye to enter and accumulate within the cells. **e,** Representative confocal images from primary human young and aged myotubes (scale bar, 50 µm) exposed to FM1 -43 fluorescent dye (green) during 4 minutes. **f,** FM1-43 intracellularly in young and aged myotubes after the inhibition of ASM by exposition to DPA. n=29-31 myotubes. Data are presented as mean + SD. ***p<0.005.

### Multi-omics integration reveals that coordinated lysosomal sphingolipid enzyme secretion reshapes intracellular lipid in aged myotubes

Given the central role of lysosomal ASM in sphingolipid metabolism and the coordinated upregulation of lysosomal secretion pathways in aged myotubes, we performed untargeted high-resolution lipidomic analysis using liquid chromatography mass spectrometry analysis of both myotubes and conditioned media (Table S3-S4). Using the supervised multi-omics integration framework DIABLO (Singh et al. 2019), we identified a combination of proteins and lipids that collectively discriminate young from aged myotubes (Fig. 4a). Multi-omics lipid integration revealed that aged myotubes accumulated simpler phospholipid species (PL18.2 and PL18.3; containing 18 carbons and 2 or 3 double bounds), at the expense of more complex phospholipids (PL20.4; containing 20 carbons and 4 double bounds) (Fig. 4a). In addition, increased secretion of peroxiderodin 6 (PRDX6) was associated with reduced intracellular PRDX6 levels (Fig. S4a-b). PRDX6 is a multifunctional enzyme with peroxidase activity toward phospholipid hydroperoxides and Ca²⁺-independent phospholipase A2 (iPLA2) activity, enabling fatty acid hydrolysis from membrane phospholipids and the conversion of lysophosphatidylcholine (LPC) into phosphatidylcholine (PC) (Lagal et al. 2024). In line with that during aging, when the specific lipid species were analyzed, a decrease of LPC was associated to an increase of PC in the conditioned media from aged myotubes (Fig. 4b). GO analysis further indicated enrichment of plasma membrane-related pathways in aged myotubes (Fig. S4c). Together, these results suggest that aging drives extensive plasma membrane lipid remodeling in skeletal muscle cells.

**Fig 4:**
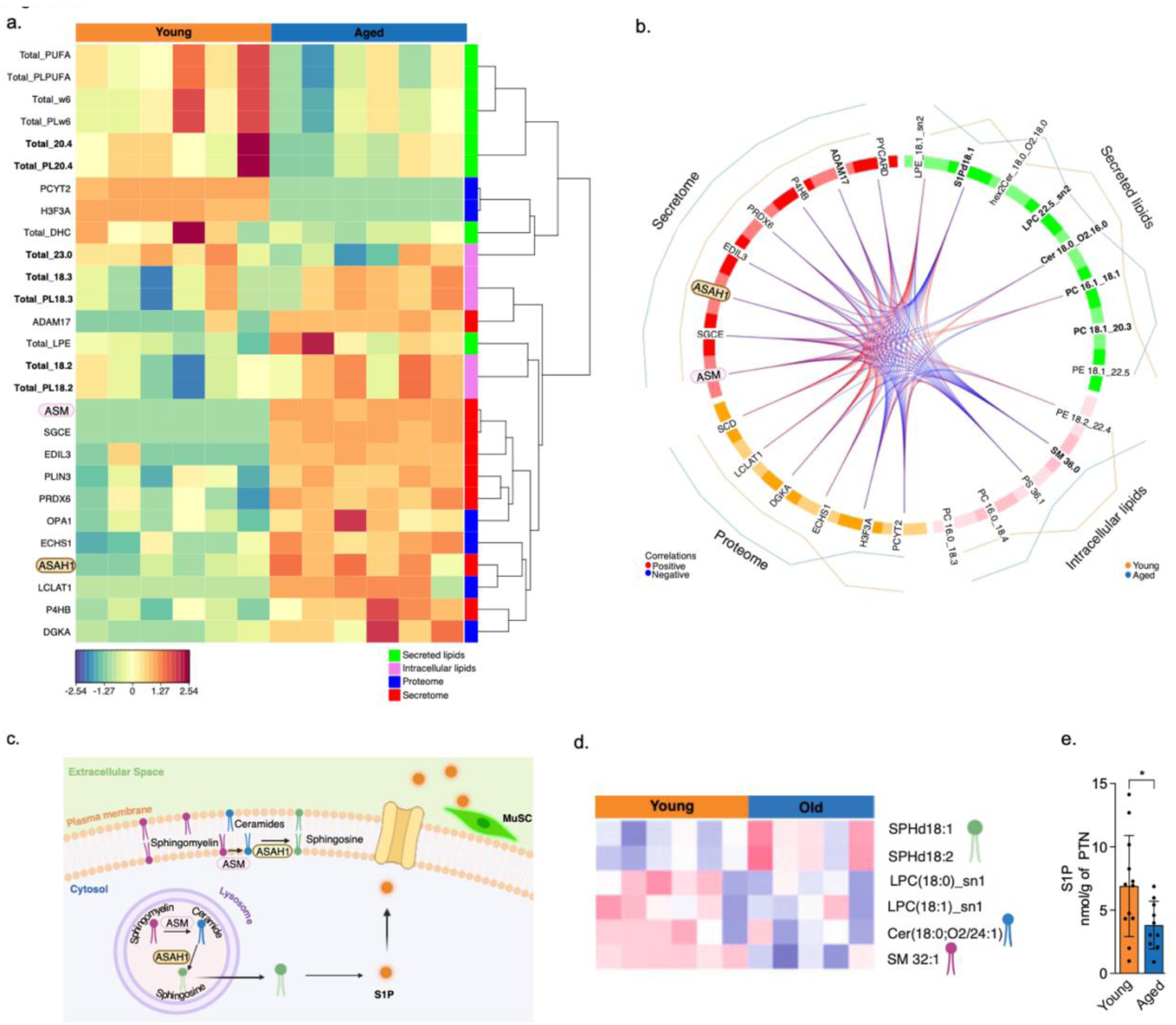
Integrative multi-Omics integration captures the age effects of sphingolipid secretion. **a**,Heatmap of normalized lipid and protein abundances in the muscle cells and conditioned media. Rows are features, and columns are samples. Color scale represents z-scored abundance (blue = low, orange=high), n=6. **b,** Circos plot for the correlations between altered intracellular and secreted proteins and lipid species identified in young and aged samples. Protein classes from the secretome and proteome are displayed on the left side of the plot. Secreted and intracellular lipid species are shown on the right. Connecting lines represent significant correlations between proteins and lipids, with red lines indicating positive correlations and blue lines indicating negative correlations. The outer lines represent the distribution profiles for young and aged groups. **c,** Schematic representation of sphingolipid metabolism at the cell membrane and in the lysosome. Sphingomyelin is converted into ceramides by ASM. Ceramide can subsequently be degraded into sphingosine by acid ceramidase (ASAH1). Sphingosine can be phosphorylated by sphingosine to form sphingosine-1-phosphate (S1P), which can exert autocrine or paracrine function in skeletal muscle. **d,** Heatmap of normalized sphingolipids abundances in myotubes from young and aged donors. Row represents sphingolipids and columns are samples. Color scale represents z-scored abundance (blue = low, red=high). n=5-6. **e,** S1P secretion levels by young and aged myotubes. n=10-12. Data are presented as mean ± s.d.. *p<0.05 vs young.

To further dissect how this lysosome-associated lipid remodeling impacts sphingolipid metabolism, we next focused on the sphingolipid axis. The age-related increase in ASM secretion (Fig. 2c and 4a) was not accompanied by changes in intracellular ASM protein levels (Fig. S4d). Instead, when focusing on the lipid species present in the conditioned media and intracellular, increased ASM secretion correlated with reduced sphingomyelin levels (SM 36.0) in aged myotubes and, unexpectedly, with a lower ceramide content (Cer 18.0_O2.16.0) in the conditioned media (Fig. 4b). This apparent paradox could be explained by the concomitant over-secretion of acid ceramidase N-acylsphingosine amidohydrolase 1 (ASAH1) (Fig. 4a-b and S4e), a lysosomal enzyme that degrades ceramides into sphingosine and free fatty acid (Fig. 4c). ASAH1 protein levels within myotubes were unchanged with age (Fig. S4f). Both ASM and ASAH1 are lysosomal enzymes whose dysregulation has been linked to human aging (Moaddel et al. 2021), further supporting the physiological relevance of this coordinated secretory response. Consistent with increased ASAH1 activity, sphingosine accumulated in aged myotubes (Fig. 4d). Sphingosine can be phosphorylated into Sphingosine-1-phosphate (S1P), a bioactive lipid that promotes cell proliferation, survival and MuSC activation (Nagata et al. 2006). Notably, despite sphingosine accumulation, aged myotubes exhibited a significant reduction in S1P secretion (Fig. 4b and 4e), indicating an age-associated uncoupling between sphingosine availability and S1P production.

### Sphingosine-1-phosphate restores the muscle stem cell reserve pool in aged muscle cultures

The age-related decline in muscle regenerative capacity has been attributed both to a reduction in the MuSC pool (Renault et al. 2002) and to alterations in the surrounding niche (Conboy et al. 2005). Exposure of aged MuSCs to a young environment has previously been shown to restore regenerative potential (Conboy et al. 2005), highlighting the importance of niche-derived signals. In our system, aged myotubes-derived niche presents increased lysosomal proteins secretion, which preserves membrane integrity while concomitantly reshaping sphingolipid signaling and reducing the PAX7+ MuSCs reserve pool. Given the reduced number of PAX7+ cells (Fig. 1d) and the decreased S1P secretion in aged cultures (Fig. 4e), we hypothesized that impaired S1P signaling contributes to MuSC pool depletion. Treatment of proliferating myoblast with 1μM S1P for 24h (Fig. 5a) significantly increased proliferation in aged cultures (Fig. 5b), but not in young cultures (Fig. 5c and Fig. S5a). Next, we assessed whether S1P could restore the reserve cell pool. Treatment of two-day differentiating aged cultures with 10μM S1P for 24h (Fig. 5d) resulted in a 2.7% increase in PAX7+ cells (Fig. 5e) and a reduction in the fusion index (Fig. 5f). In contrast, S1P had no effect on reserve cell number or differentiation in young cultures (Fig. 5g-h and Fig. S5b). Overall, S1P supplementation restored the PAX7+ reserve stem cell pool in aged cultures, partially rescuing levels observed in young conditions.

**Fig 5:**
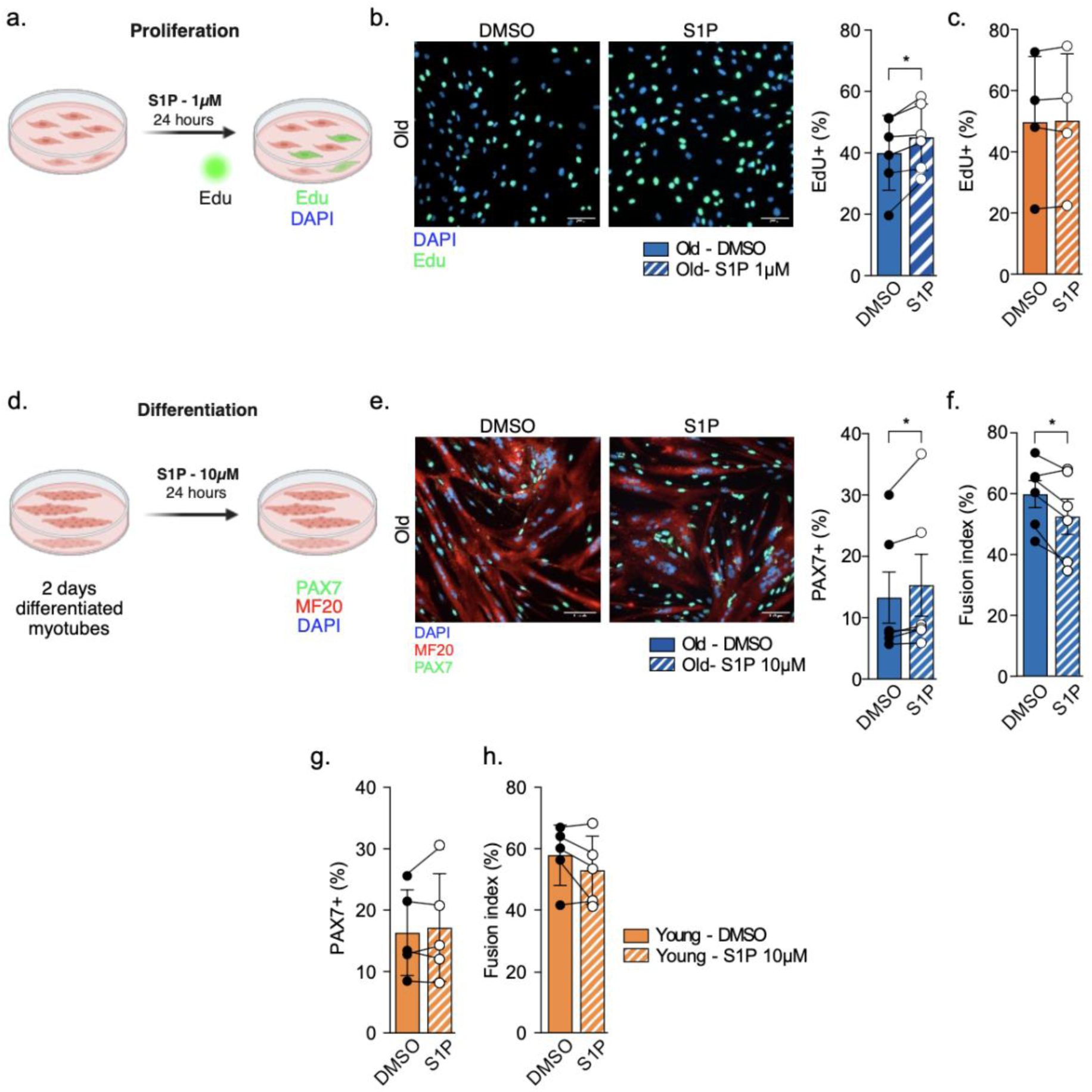
S1P treatment effects in young and aged cultures. **a**,Schematic representation of the experimental setup. Myoblasts were treated with S1P, 1µM, for 24 hours prior to Edu analysis. **b,** Edu+ analysis in aged myoblast after S1P 1µM treatment, n=6 cell lines, **c,** EdU+ analysis in young myoblast after S1P 1µM treatment, n=4 cell lines. **d,** Schematic representation of the experimental setup. Myotubes differentiated for 2 days were treated with S1P, 10µM for 24 hours prior to analysis. **e,** MuSCs immunostaining in aged myotubes after S1P 10µM treatment (scale bar, 100 <u>µ</u>m). PAX7 (green), MF20 (red) and DAPI (blue) and PAX7 positive cells in the aged culture treated with S1P 10µM, n=6 cell lines. **f,** Fusion index analysis in aged myotubes treated with S1P 10µM, n=6 cell lines**. g,** PAX7 positive cells in the young culture treated with S1P 10µM, n=4 cell lines. **h,** Fusion index analysis in young myotubes treated with S1P 10µM, n=4 cell lines. Data are presented as mean ± s.d. *p<0.05 vs DMSO.

## Discussion

A growing body of evidence indicates that the regenerative decline of aged skeletal muscle is influenced by both intrinsic MuSC defects and changes in secreted factors from the surrounding environment (Conboy et al. 2005). In addition to its contractile function, skeletal muscle acts as a secretory organ, releasing a wide range of proteins, RNAs, and metabolites (Pedersen & Febbraio 2012). The muscle secretome regulates tissue homeostasis through autocrine and paracrine mechanisms (Pedersen & Febbraio 2012), and modulates MuSC behavior, including quiescence, activation, and self-renewal, thereby contributing to the maintenance of the stem cell pool (Yin et al. 2013). Importantly, systemic factors also play a key role in muscle aging. Heterochronic parabiosis experiments demonstrated that exposure of aged animals to a young systemic environment improves MuSC function, whereas young animals exposed to aged circulation exhibit impaired regenerative capacity (Conboy et al. 2005). However, these approaches do not establish skeletal muscle as the source of the circulating factors involved, highlighting the need to directly define muscle-derived secreted factors and their age-associated alterations. Here we show that aging remodels the muscle secretome and that dysregulation of sphingolipid metabolism directly impairs MuSC pool and regenerative capacity.

The ASM–ceramide system is a well-established regulator of inflammatory responses, notably by controlling the release of pro-inflammatory cytokines (Grassmé et al. 2008). In the skeletal muscle, ASM activity is induced following injury (Roux-Biejat et al. 2021) and in pathological contexts associated with membrane instability (Defour et al. 2014), where it contributes to membrane repair through lysosomal exocytosis and compensatory endocytosis (Tam et al. 2010). Treatment of muscle cells from a patient with dysferlinopathy (a myopathy characterized by membrane damage) with recombinant ASM, improves plasma membrane repair (Bittel et al. 2022). Consistent with these roles, our results suggest that increased ASM secretion during aging contributes to preservation of membrane integrity, as its inhibition leads to membrane damage. These observations indicate that elevated ASM secretion during aging represents an adaptive response to membrane stress. Moreover, increased circulating ASM levels and lysosomal dysfunction have been reported during aging (Park et al. 2018), suggesting that age-associated alterations in muscle secretion may also contribute to systemic alterations. Lipid composition is a critical determinant of plasma membrane organization and cellular function (Lagal et al. 2024). Our multi-omics analyses reveal that aging profoundly remodels lipid composition in human skeletal muscle cells, with potential consequences for both membrane integrity and signaling. In particular, aged myotubes exhibit reduced sphingomyelin levels, consistent with altered membrane organization. This is in line with previous observations that disruption of lipid homeostasis, including through loss of PRDX6 function, destabilizes membrane lipid rafts through reduced sphingomyelin content (Lagal et al. 2024). In parallel, the accumulation of sphingosine in aged myotubes may further contribute to cell membrane alterations, by promoting biophysical changes in cellular membranes (Zupancic et al. 2014). Additionally, it has been described an age-related reduction of the plasma levels of polyunsaturated fatty acids (PUFAs) in females (Poliezhaieva et al. 2026), as our data demonstrated a reduced of the secretion of PUFAs by aged myotubes. The low levels of PUFA in the plasma of aged individuals were negative correlated to the performance in the walking speed test (Poliezhaieva et al. 2026). Together, these lipid alterations support a model in which increased ASM secretion drives SM hydrolysis, leading to a shift in sphingolipid balance that impacts membrane properties, representing an adaptive response to preserve membrane integrity under stress, but at the cost of altered sphingolipid balance.

Importantly, this imbalance directly impacts MuSCs regulation. We show that aging-associated secretion of ASM and ASAH1 is accompanied by reduced availability of S1P, a bioactive sphingolipid known to promote MuSC activation (Nagata et al. 2006) and proliferation (Calise et al. 2012). S1P can be secreted by muscle cells, including via extracellular vesicles (EVs), where it exerts biological activity (Hakkar et al. 2025). Restoration of S1P levels through exogenous treatment was sufficient to rescue MuSC proliferation and partially restore the reserve cell pool in aged cultures, functionally linking altered sphingolipid metabolism to MuSC dysfunction. Although not directly assessed here, altered activity of sphingosine kinase 1 (SPHK1), which converts sphingosine into S1P, may contribute to the reduced S1P availability observed during aging, consistent with previous work highlighting a role for SPHK1/S1P signaling in skeletal muscle regeneration (Sassoli et al. 2011).

Some limitations of this study should be considered. First, our analyses were performed using *in vitro* differentiated human muscle cells, which do not fully capture the complexity of the *in vivo* niche, including interactions with immune, vascular and neural compartments. Although we identify ASM and S1P as key regulators of MuSC behavior, the relative contribution of muscle-derived versus circulating sphingolipids remains unclear. Second, while our functional assays support a causal role for sphingolipid imbalance in regulating the MuSC pool, *in vivo* validation will be required to determine the impact of ASM and S1P modulation on muscle regeneration. Finally, the broader contribution of other age-associated secreted factors identified in our proteomic dataset remains to be explored.

Together, our findings support a model in which aging-associated remodeling of the muscle secretome disrupts sphingolipid homeostasis, preserving membrane integrity through increased ASM-dependent sphingomyelin hydrolysis while concurrently limiting S1P availability, thereby impairing MuSC maintenance. These results identify sphingolipid metabolism as a mechanistic link between muscle aging and stem cell dysfunction, and highlight ASM and S1P as potential targets to preserve regenerative function in aged muscle. These findings may have broader implications for other aging tissues in which lipid-mediated intercellular communication regulates stem cell function.

## Materials and Methods

### Human muscle cells

The cells were obtained from mechanically dissociated human muscles biopsies as previously described (Bigot et al. 2009). Muscle biopsies were obtained from the Myobank-AFM (bank of tissues for research, a partner in the EU network EuroBioBank) in accordance with European recommendations and French legislation. Additional cell lines were obtained in collaboration with the University of Kentucky Center for Muscle Biology as previously described (Walton et al. 2019). For muscle and age information from the donors see Supplementary figure 6.

The myogenic cell population was enriched using a CD56 magnetic bead sorting system (MACS, Miltenyi Biotech). The myoblasts were cultured in proliferation medium (1 volume of M199 (ThermoFisher, 41150020), 4 volumes of Dulbecco’s modified Eagle’s medium (DMEM) (ThermoFisher; 61965-026), 20% fetal bovine serum). For the myotubes studies, cells were differentiated during 3 days in DMEM.

*Generation of cell proteome*.

Cell pellets of these same cultures have been harvested for proteomic analyses. Each pellet was lysed in a buffer containing 200 mM triethylammonium bicarbonate (TEAB), 2% SDS, 50 mM CAA (chloroacetamide) and 10 mM tris(2-carboxyethyl) phosphine (TCEP), then heated for 5 min at 95°C. After gel assay with RPMI lysate as a reference, 30 micrograms of each pellet were digested using trypsin (Promega) and S-Trap Micro Spin Column was used according to the manufacturer’s protocol (Protifi, Farmingdale, NY, USA). Peptides were then speed-vacuum dryed, then were solubilized in 2% trifluoroacetic acid (TFA) and fractionated by strong cationic exchange (SCX) Stage-Tips, as described(Kulak et al. 2014). After drying, peptides from SCX Stage-Tip, the 5 fractions were solubilized in 10 μL of 0.1% TFA containing 10 % acetonitrile (ACN). One µL of peptides were loaded and separated by Reverse phase C18 using an u3000 RSLC chromatographer from Dionex on a 2 µm particle size, 100 Angström pore size, 75 µm internal diameter, 25 cm length C18 reverse phase analytical column with a 3h binary gradient from 99% solution A (0.1% formic acid in H2O) to 55% solution B (80% ACN, 0.085% formic acid) before injection into an Orbitrap Fusion Mass Spectrometer MS (Thermo Scientific). MS data acquisition was performed throughout the elution process in a data-dependent scheme (top speed mode in 3 seconds) with full MS scans acquired with the orbitrap detector, followed by HCD peptide fragmentation and Ion trap fragment detection of the most abundant ions detected in the MS scan. Mass spectrometer settings for full scan MS were: 1.0E6 AGC, 60,000 target resolution, 350-1500 m/z range, maximum ion injection time (MIIT) of 60 ms. HCD MS/MS fragmentation was permitted for 2-7+ precursor ions reaching more than 5.0E3 minimum intensity. Quadrupole-filtered precursors within 1.6 m/z isolation window were fragmented with a Normalised Collision Energy setting at 30. 1.0E5 AGC Target and 60 ms MIIT were the limiting ions accumulation values. The Ion-trap detector was used for its fast and sensitive detection capabilities. A 30 seconds dynamic exclusion time was set. Maxquant software version 2.0.3.0(Cox et al. 2014) was used for the data extraction from the raw MS files from mass spectrometer. The databases used were the human sequences from the Uniprot database (March 2020) and a list of in-house frequent contaminant sequences. The cleavage specificity was trypsin’s with maximum 2 missed cleavages. Carbamidomethylation of cysteines was set as constant modification, whereas acetylation of the protein N terminus and oxidation of methionines were set as variable modifications. The false discovery rate was kept below 1% on both peptides and proteins. The normalization was permitted and “match between runs” (MBR) option was allowed with a match time window of 1 min and an alignment time window of 30 min.

### Proliferation analysis

For the proliferation assay, 2500 cells were plated on a µ-Slide 8 Well (Ibidi, Gräfelfing, Germany) in proliferation media and allowed to attach in a 5% CO_2_ incubator at 37°C during 24 hours. Cells were then incubated with 10µM of EdU for 14 hours. Following the cells were washed with 1x PBS and fixed with 3.6% of Paraformaldehyde (PFA). Fixed cells were washed with 1x PBS and stored at 4°C for subsequent immunofluorescence using the *Click-iT® EdU Alexa Fluor® 488* Imaging Kit (Invitrogen; C10337). For nuclear staining, the slides were incubated with DAPI for 5 min, rinsed 2 times and mounted with mounting media (DAKO). Images were acquired using Zeiss Axio Observer Apoptome. The number of divisions were calculated as previously described (Bigot et al. 2015).

### Extracellular flux (XF) analysis

Between 12,000-13,000 cells were plated in a Seahorse XF96 Cell culture microplate coated with 0,5% gelatin and allowed to attached for 24 hours. Following cells were washed with 1x PBS and incubated with differentiation media (DMEM, high glucose GlutaMax supplemented with 4% heat-inactivated horse serum, 1% 100 mM sodium pyruvate)(Sapaly n.d.) during 3 days. The mitochondrial oxygen consumption rate (OCR) was performed using a Seahorse analyzer (XF96, Agilent Technologies), following the Seahorse XF96 Cell Mito Stress Test(Gu et al. 2021). Briefly, in the day of the analysis cells were rinsed with 1x PBS once and incubated with Seahorse XF RPMI medium containing 1mM pyruvate, 2mM glutamine and 10mM glucose, during 1 hour at 37 °C non-CO_2_ incubator. The OCR was measured using 1.5µM oligomycin to inhibit ATP synthase, 5µM carbonyl cyanide-p-trifluoromethoxyphenylhydrazone (FCCP) to uncouple oxidative phosphorylation (OXPHOS), and 0.5µM rotenone and antimycin to inhibit mitochondrial respiration, to obtain the non-mitochondrial oxygen consumption. Basal respiration was calculated by subtracting the non-mitochondrial OCR from the last OCR measurement before the first injection (Paradis & Technologies n.d.). ATP-production was calculated by subtracting the minimum OCR value after oligomycin injection from the basal OCR (Lima et al. 2023). Maximum respiratory was determined by subtracting the values from the non-mitochondrial OCR from the OCR values obtained after FCCP addition (Paradis & Technologies n.d.). Proton leak was calculated by subtracting the nonmitochondrial OCR values from the minimum values obtained after the oligomycin injection (Paradis & Technologies n.d.). All Seahorse measurements were normalized by protein quantified by Pierce™ BCA Protein (Thermo Fisher Scientific).

### Amyloid deposit analysis

Between 65,000-70,000 cells were plated on a 12mm glass cover slip coated with 0,5% gelatin. The following day, cells were washed with 1x PBS and incubated with DMEM (differentiation media) for 3 days. The myotubes were fixed using 3.6% formaldehyde, then permeabilized and blocked using 0.5% Triton X-100-PBS with 2% FBS for 1 hour at room temperature. Proteostat® Protein aggregation assay (Enzo Life Science, New York, USA) was performed. Cells were incubated with Proteostat® dye (1:200) and MF20 (1:20) for 2 hours at room temperature. Following, cells were washed with 1x PBS and incubated with secondary antibody anti-mouse IgG2b AlexaFluor 647, (1:400, Invitrogen™) for 1 hour at room temperature. For nuclear staining, the slides were incubated with DAPI for 5 min, rinsed 2 times and mounted with mounting media (DAKO). Images were acquired using 40x magnification at Nikon Ti-2 inverted microscope. The area of amyloid deposit (called aggresomes) was analyzed using the Fiji software(Schindelin et al. 2012) and normalized by the area of the myotubes in the image.

### Fusion Index

The fusion index was quantified using Fiji software(Schindelin et al. 2012). It was calculated as the percentage of nuclei located within MF20-positive myotubes relative to the total number of nuclei (Bensalah et al. 2022). Images were acquired from randomly selected fields and quantified in a blinded manner, with at least 500 nuclei per condition.

### PAX7 immunostaining

Between 36,000-41,000 cells were plated on a µ-Slide 8 Well (Ibidi, Gräfelfing, Germany) in proliferation media. The following day, cells were washed with 1x PBS and incubated with DMEM only (differentiation media) for 3 days. The myotubes were fixed using 3.6% formaldehyde, then permeabilized, blocked, and stained as previously described (Duguez et al. 2013). Primary antibodies used were anti-Pax7 (PAX7, 1:20, Developmental Studies Hybridoma Bank (DSHB)), anti-myosin heavy chain (MF20, 1:50, DSHB). The secondary antibodies used were goat anti-mouse IgG1 488 or anti-mouse IgG2b AlexaFluor 555, (1:400, Invitrogen™). For nuclear staining, the slides were incubated with DAPI for 5 min, rinsed twice and mounted with mounting media (DAKO). Images were acquired using Zeiss Axio Observer Apoptome. The PAX7+ cells were analyzed using the Fiji software (Schindelin et al. 2012).

### Generation of secretome

Samples of conditioned media necessary for the generation of the secretome were harvested from 3-days differentiated human myotubes. Supernatants were then centrifugated for 10min at 4°C and 300g to get rid of cellular debris; supernatant was collected and then centrifugated for 20min at 4°C and 2000*g*. Following, to concentrate the conditioned media, the supernatant was centrifuged using Amicon ®Ultra-4 columns (UFC800396) during 35min at 4000*g*.

All samples were solubilized volume by volume in lysis buffer containing 400 mM TEAB, 4% SDS, 100 mM CAA, 20 mM tris(2-carboxyethyl) phosphine (TCEP), then they were boiled 5 min at 95°C. Each sample were digested using trypsin (Promega) and S-Trap Micro Spin Column was used according to the manufacturer’s protocol (Protifi, Farmingdale, NY, USA).

Peptides were then speed-vacuum dryed, then were solubilized in 10 μL of 0.1% TFA containing 10 % acetonitrile (ACN). nLC-MS/MS analyses were performed on a Dionex U3000 RSLC nano-LC-system coupled to a TIMS-TOF Pro mass spectrometer (Bruker Daltonik GmbH, Bremen, Germany). One μL of peptides were loaded, concentrated and washed for 3min on a C18 reverse phase column (5μm particle size, 100Å pore size, 300 μm inner diameter, 0.5 cm length, from Thermo Fisher Scientific). They were separated on an Aurora C18 reverse phase resin (1.7μm particle size, 120Å pore size, 75μm inner diameter, 25cm length mounted to the Captive nano Spray Ionisation module, from Ion Opticks, Middle Camberwell Australia) with a 2h run time with a gradient ranging from 98% of solvent A containing 0.1% formic acid in milliQ-grade H2O to 35% of solvent B containing 80% acetonitrile, 0.085% formic acid in mQH_2_O.

Some samples clogged the column, so the peptides were dried and taken up in 50µL of TFA 2%. In a P200 cone without filter, after cutting the top of the filter, 6 layers of SCX membranes (3M empore: solid phase extraction disks) were added(Rappsilber et al. 2007). The SCX phase was activated with 100µl of ACN 100%, then washed twice with TFA 0.1%. The sample was passed over the SCX phase, and the phase was washed a first time with 100µl of TFA 0.5% and a second time with 100 µL of TFA 0.5% + ACN 50%. Peptide elution was then performed three time with 20µL of NH4OH 5% + ACN 80%. Peptides were dried then solubilized in 10 μL of 0.1% TFA containing 10 % ACN. One µl of peptide was loaded onto the same column as described previously but with a 1h gradient going from 98% of solvent A to 10% of solvent B in 14 min, then 20% of solvent B in 25 min and 40% of solvent B in 38 min.

The mass spectrometer acquired data throughout the elution process and operated in DDA PASEF mode with a 1.89 second/cycle, with Timed Ion Mobility Spectrometry (TIMS) mode enabled and a data-dependent scheme with full MS scans in PASEF mode. This enabled a recurrent loop analysis of a maximum of the 120 most intense nLC-eluting peptides which were CID-fragmented between each full scan every 1.89 second. Ion accumulation and ramp time in the dual TIMS analyser were set to 166 ms each and the ion mobility range was set from 1/K0 = 0.6 Vs cm^−2^ to 1.6 Vs cm^−2^. Precursor ions for MS/MS analysis were isolated in positive mode with the PASEF mode set to « on » in the 100-1.700 m/z range by synchronizing quadrupole switching events with the precursor elution profile from the TIMS device. Precursors for MS/MS were picked from an intensity threshold of 1000 arbitrary units (a.u.) and re-sequenced until reaching a ‘target value’ of 20.000 a.u. considering a dynamic exclusion of 0.40 s elution gap.

The mass spectrometry data were analysed using Maxquant version 1.6.17.0(Tyanova et al. 2015). The database used for in silico generation of spectral library was a concatenation of Human sequences from the Swissprot database (release 2020-03) and a list of contaminant sequences from Maxquant and from the cRAP (common Repository of Adventitious Proteins). The enzyme specificity was that for trypsin. Carbamidomethylation of cysteines was set as constant modification, whereas acetylation of the protein N terminus and oxidation of methionines were set as variable modifications. The false discovery rate was kept below 1% on both peptides and proteins. The “match between runs” (MBR) option was allowed.

### Lipidomics analysis

Conditioned media for the lipidomic analysis was collected and concentrated following the same protocol described in *“Generation of secretome”*. The lipidomic analysis from the cell pellet was performed using the lipid extract from cells harvested after 3 days of differentiation. The lipids were extracted according to a modified Folch method(Folch et al. 1957). To analyze secreted lipids, 50µl of conditioned media were supplemented with internal standards. To the quantification of intracellular lipids, the cell pellets were resuspended in water (1,000,000 cells/μl) and 50µl of cell homogenate were supplemented with internal standard. Following, the lipids were extracted using 1.5ml of chloroform and 750µl of methanol and vortexed for 1 min. The phase separation was trigged by addition of 450µl of ammonium carbonate 250mM and vortexed for 1 min. The extracted lipids were dried and resuspended in 50µl of LC/MS solvent. The lipids were quantified by LC-ESI/MS/MS using a Prominence UFLC and a QTrap 4000 mass spectrometer. Samples (4μl) were injected into a Kinetex. HILIC 2.6µm 2.1×150mm column (Phenomenex, CA, USA). Mobile phase consisted of water and acetonitrile containing ammonium acetate and acetic acid. Lipid species were detected using schedule multiple reaction monitoring (sMRM) in the positive-ion mode for sphingolipids, LPC, LPE, PA and PS reflecting the headgroup fragmentation of each lipid class and in the negative ion mode for PC, PE, PI, PG reflecting fatty acid chain fragmentation. An in-house developed R script was used to correct isotopic contribution on MRM signals from HILIC injections. Lipid features whose variability exceed 30% in the QC were removed.

### S1P and sphingosine analysis

100µl of concentrated conditioned media or cell homogenate prepared from the cell pellet were supplemented with deuterated S1P, 180µl of 1% formic acid and 680µL of methanol. Samples were vortexed for 1 min and centrifuged at 16,000*x g* for 2 minutes. The supernatants were transferred to LC vials and dried, and sphingosines were resuspended in 50μL of 95% methanol. The lipids were quantified by LC-ESI/MS/MS using a Prominence UFLC and a QTrap 4000 mass spectrometer. Samples (4μl) were injected to an Atlantis HILIC 2.6µm 2.1×50mm column (Waters Corporation, MA, USA). Mobile phase consisted of water containing 50mM ammonium formate and 0.2% formic acid and acetonitrile containing 0.2% formic acid. Lipid species were detected using scheduled multiple reaction monitoring (sMRM) in the positive-ion mode. An in-house developed R script was used to correct isotopic contribution on MRM signals from HILIC injections. Lipid features whose variability exceed 30% in the QC were removed.

### ASM treatment

Between 36,000-41,000 cells were plated on a µ-Slide 8 Well (Ibidi*®*) in proliferation media. The following day, the cells were washed with 1x PBS and incubated with DMEM (differentiation media). After two days in differentiation media, the myotubes were treated with human recombinant ASM protein (R&D Systems, 5348-PD-010, Lot: RUB0221121) 6U/L, as previously described(Bittel et al. 2022), for 24 hours. The myotubes were fixed using 3.6% formaldehyde, then permeabilized, blocked, and stained as previously described(Duguez et al. 2013) for subsequent analysis.

### S1P treatment

For the analysis of S1P effects during the differentiation, between 36,000-41,000 cells were plated on a µ-Slide 8 Well (Ibidi*®*) in proliferation media. After two days in differentiation media, the myotubes were treated with 10µM of S1P (Thermo Scientific), as previously described(Nagata et al. 2006), for 24 hours. For the analysis of the S1P effects during proliferation, 25,000 cells were plated on a µ-Slide 8 Well (Ibidi*®*) in proliferation media. The following day, the myoblasts were treated with 1µM of S1P during 24 hours as previously described (Calise et al. 2012). The cells were fixed using 3.6% formaldehyde, then permeabilized, blocked, and maintained at 4°C for subsequent analysis.

### ASM inhibition

3 days differentiated myotubes on µ-Slide 8 Well (Ibidi*®*) were incubated during 60 minutes with 30µM desipramine (DPA) (Sigma-Aldrich) at 37 °C, 5% CO_2_, as previously described(Bittel et al. 2022).

### Live imaging of FM1-43 influx

To induce cell membrane damage 3 days differentiated myotubes were pre-incubated with the bacterial pore-forming toxin, Streptolysin O (SLO) (Sigma-Aldrich) 600 ng/ml during 5 minutes at 4 °C (Tam et al. 2010). Following cells were incubated with FM1-43 (Invitrogen) 5µM (Carmeille et al. 2017), with or without Streptolysin O (SLO) (Sigma-Aldrich), and imaged during 4 minutes as previously described (Tam et al. 2010), at 1 frame/10s using a 40x magnification at Nikon Ti-2 inverted microscope. Quantitative analysis of intracellular fluorescence was performed using the Fiji software(Schindelin et al. 2012).

### Statistical Analysis

No statistical methods were used to predetermine sample size. Statistical analysis was performed using GraphPad Prism (version 10.4.1). Data normality was assessed using the Shapiro-Wilk test. Differences between young and aged groups were assessed using unpaired two-tailed Student’s *t* test for normally distributed data and Mann-Whitney *U* test for non-normally distributed data. Comparison between treated and control groups were performed using paired two-tailed Student’s *t* test. Analysis of the OCR time course comparing young and aged groups was conducted using two-way ANOVA followed by Tukey’s multiple-comparison test. Sample sizes (*n)* and exact *p* values are reported in the corresponding figures.

### Statistical Analysis of Label-Free Proteomics Data

Label free quantification (LFQ) intensity values derived from mass spectrometry were log2-transformed to stabilize variance and approximate a normal distribution. Proteins annotated as contaminants, reverse database hits, or identified only by site were removed prior to analysis. To ensure robust quantification, proteins were retained only if they were quantified in at least 2 biological replicates from at least one experimental group. To account for technical variability across samples, LFQ intensities were median normalized. Comparisons of protein abundance between experimental groups were performed using analysis of variance (ANOVA), to allow the simultaneous evaluation of the main effects of age and sex, as well as their potential interaction. Proteins exhibiting statistically significant differences across groups were identified based on the selected significance threshold, with multiple testing correction, (e.g., *padj* < 0.05).

### Bioinformatic analysis

In order to explore the sources of variation in the data, Centered Principal Component Analysis (PCA) was performed on the 500 most variable proteins. From the list of differentially detected proteins, enrichment of known biological functions was detected using Over Representation Analysis (ORA)(Boyle et al. 2004) using all expressed genes as universe. To detect more subtle enrichment signals, Gene Set Enrichment Analysis (GSEA) (Subramanian et al. 2005) was performed on the entire gene set, with a minimum size of gene set accepted of 10. All collections available in the Molecular Signatures Database (http://www.gsea-msigdb.org/gsea/msigdb/collections.jsp) were queried. Redundant terms were manually removed, and most interesting ones according to their biological activity were further studied.

### Integrative multi-omics analysis

Multi-omics data integration was performed using the mixOmics R package (version 6.6.2). We applied the DIABLO (Data Integration Analysis for Biomarker discovery using Latent cOmponents) framework (Singh et al. 2019), a supervised multivariate method designed to identify correlated and discriminative variables across multiple omics datasets. Four data blocks were integrated, namely, Secretome, Lipids, Cellular markers, and Lipid species derived from cell pellets. Integration was conducted using sparse Partial Least Squares Discriminant Analysis (sPLS-DA) within the DIABLO framework to maximize covariance between latent components while selecting the most discriminative features separating “Young” and “Aged” subjects. Feature selection was performed using sparsity constraints to retain the most informative variables per block. The first latent component (Component 1) was used to evaluate cross-block associations and group discrimination. Statistical significance of classification performance was evaluated using the Wilcoxon test.

## Acknowledgements

We thank Virginie Salnot and Philippe Chafey from Proteom’IC core facility of Institut Cochin in Paris for data acquisition and analyses of the proteomic data. We thank the MyoLine platform and the MyoImage platform of the Center of Research in Myology as well as the Myobank-AFM biobank from the Institute of Myology. We thank Kamel Mamchaoui for the cell culture training. We thank Charlotte A. Peterson and the University of Kentucky Center for Muscle Biology for providing human muscle cells.

## Author Contributions

Conceptualization and experimental design: BC, CT, AB ; Data acquisition and analyses: BC, NP, MB, JO, ML, MP, JB, MK, LGS; Figure composition and editing: BC, CT, AB; Original draft: BC, ABo, VM, GBB, CT, AB; Writing – Review & Editing: BC, BP, MB, JO, ML, MP, JB, MK, LGS, EN, ABo, VM, GBB, CT, AB Supervision: EN, CT, AB ; Funding Acquisition: EN, ABo, VM, GBB, CT, AB. All authors have reviewed and approved the final manuscript for publication.

## Data Availability Statement

The data that supports the findings of this study are available in the supplementary material of this article.

