## Supplementary figures for "Aging remodels the human muscle secretome, impairing stem cell via sphingolipid metabolism"

Figure S1.

a.

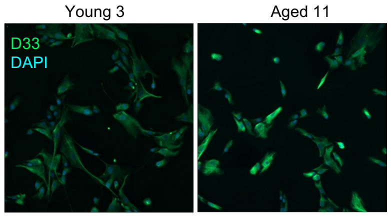


b.

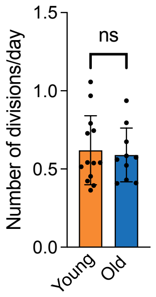


c.

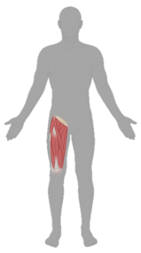

| | Age | Muscle | Sex |
|---|---|---|---|
| Young 1 | 15 | Quadriceps | F |
| Young 2 | 17 | Quadriceps | F |
| Young 3 | 19 | Quadriceps | F |
| Young 4 | 24 | Quadriceps | M |
| Young 5 | 21 | Quadriceps | M |
| Young 6 | 15 | Gluteo Maximus | F |
| Old 1 | 86 | Fascia Lata | F |
| Old 2 | 74 | Quadriceps | F |
| Old 3 | 80 | Quadriceps | M |
| Old 4 | 60 | Fascia Lata | M |
| Old 5 | 79 | Quadriceps | M |
| Old 6 | 76 | Quadriceps | F |

d.

| | Total POI | Up in old | Down in old |
|---|---|---|---|
| Proteome | 408 | 186 | 222 |
| Secretome | 127 | 102 | 25 |

e.

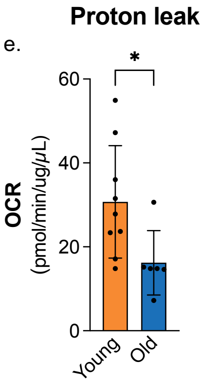


**a,** Desmin (D33) immunofluorescence staining in young and aged myoblasts. b, Number of divisions per day from young and aged donors myoblast. n=10-13. Data is presented as mean ± s.d. **c,** Table with the age, skeletal muscle source, and sex of young and aged donors included in the proteomics study. **d,** Table with the number of proteins of interest (POI) in the proteome and secretome analysis between young and aged donors. **e,** Proton leak analysis during a mitochondrial stress test, n=7-9. Data is presented as mean +SEM.

Figure S2.

a.

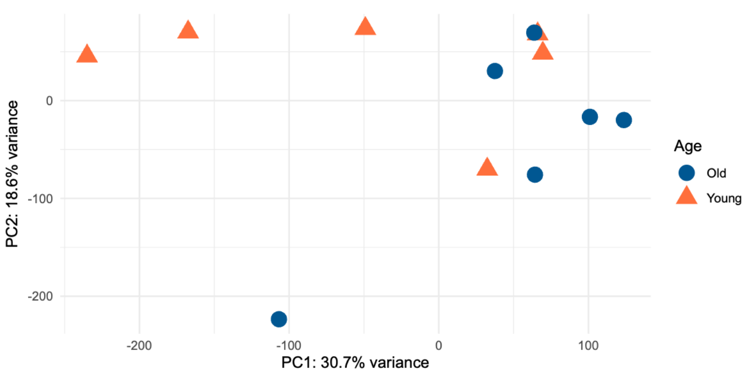


**a,** PCA analysis from the secretome of young and aged donors myotubes. n=6.

Figure S3.

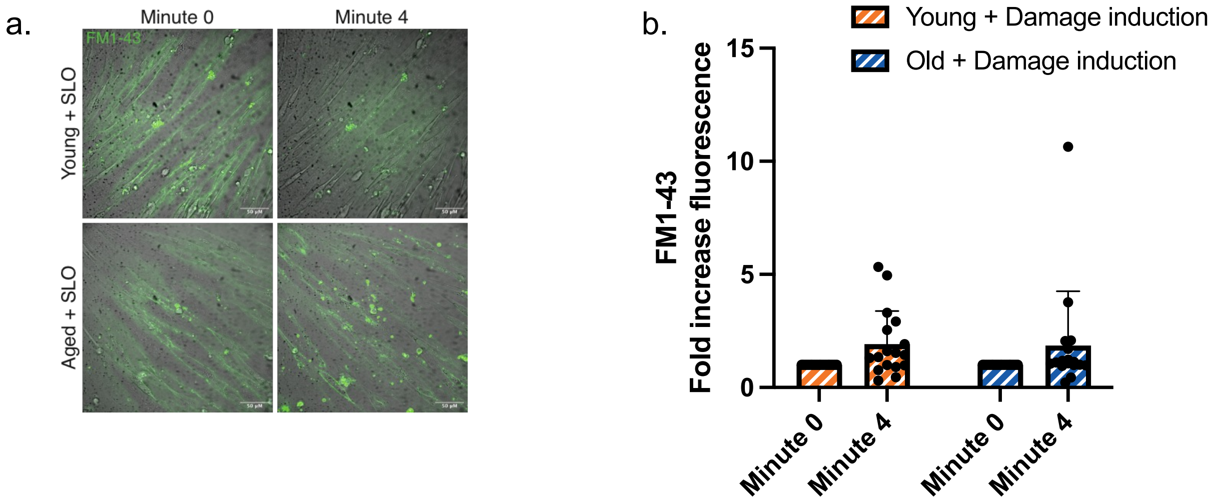


**a,** Representative confocal images from primary human young and aged myotubes (scale bar, 50 µm) exposed to FM1-43 fluorescent dye (green) and to, streptolysin-O (SLO) during 4 minutes. **b,** FM1-43 intracellularly in young and aged myotubes after the exposure to the bacterial pore forming toxin, streptolysin-O (SLO). n=29-31 myotubes. Data is presented as mean + SD. ***$p$<0.005.

Figure S4.

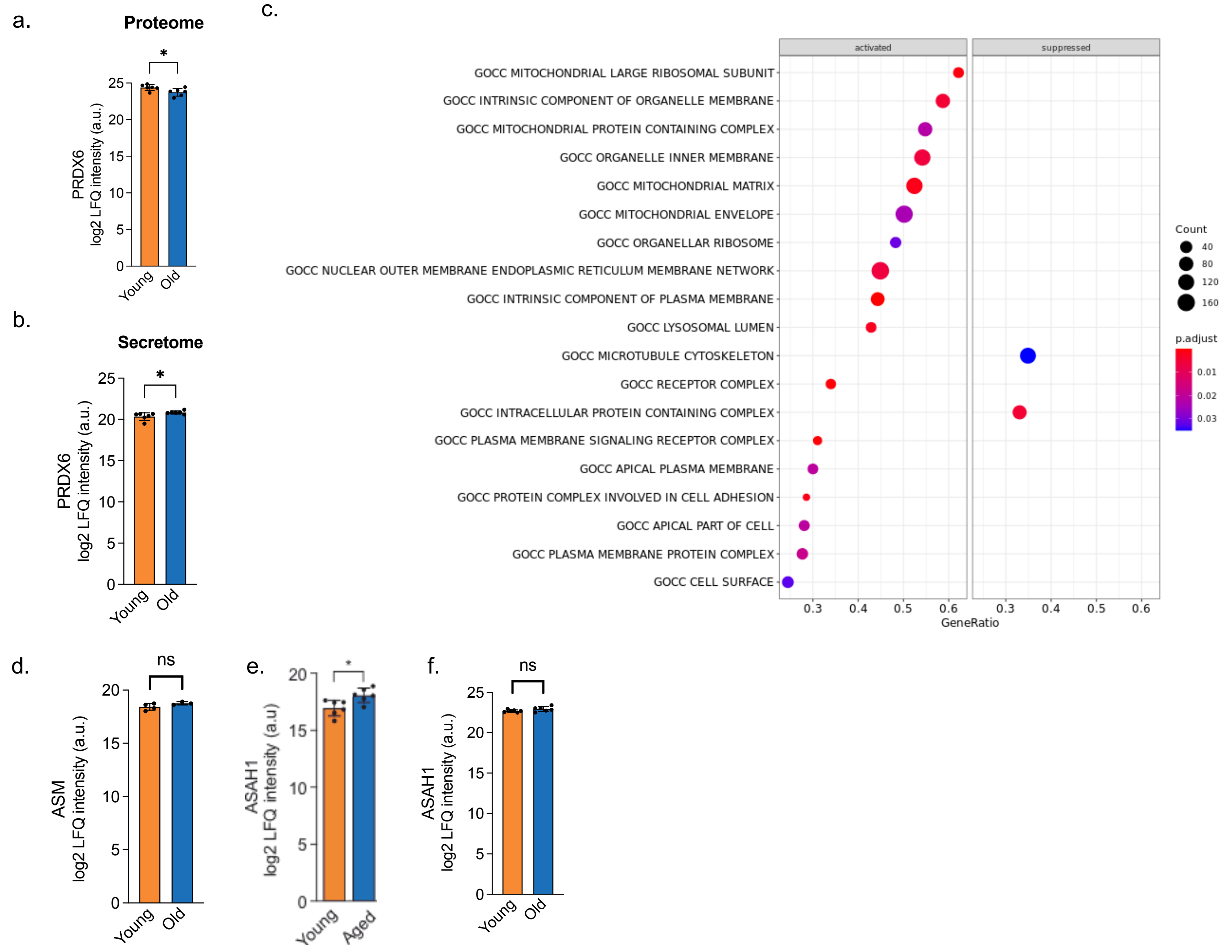


**a,** PRDX6 protein levels in the young and aged myotubes. n=6. **b,** PRDX6 protein levels in the conditioned media from young and aged myotubes. n=6. Data are presented as mean ± s.d. **p*<0.005.. **c,** Enrichment analysis of proteomic data using the "Cell Composition" from the Gene ontology database, comparing young and aged myotubes. **d,** ASM intracellular protein content. Data is presented as mean ± s.d. **e,** ASAH1 protein levels in the conditioned media from young and aged myotubes. n=6. Data is presented as mean ± s.d. **f,** ASAH1 intracellular protein content. n=6. Data is presented as mean ± s.d.

Figure S5.

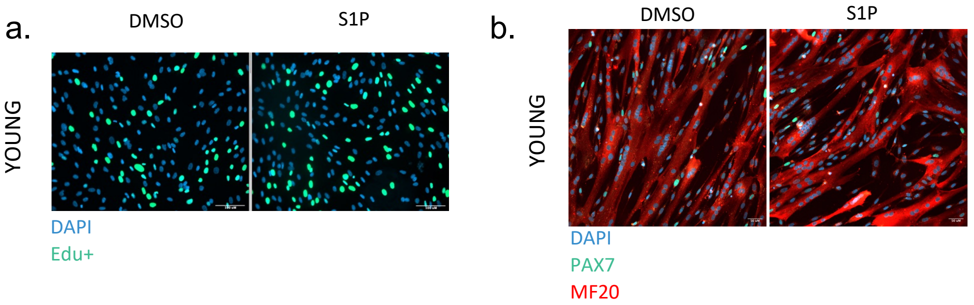


**a,** Representative image from the Edu+ analysis in young myoblast after S1P 1μM treatment. **b,** Representative image from the immunofluorescence staining for PAX7 and MF20 in the young myotubes culture.

Figure S6

a.

| | Age | Muscle | Sex |
|---|---|---|---|
| Young 1 | 15 | Quadriceps | F |
| Young 2 | 17 | Quadriceps | F |
| Young 3 | 19 | Quadriceps | F |
| Young 4 | 24 | Quadriceps | M |
| Young 5 | 21 | Quadriceps | M |
| Young 6 | 15 | Gluteo Maximus | F |
| Young 7 | 26 | Quadriceps | M |
| Young 8 | 25 | Quadriceps | F |
| Young 9 | 15 | Latissimus dorsi | F |
| Young 10 | 20 | Quadriceps | M |
| Young 11 | 17 | Quadriceps | M |
| Young 12 | 20 | Fascia Lata | M |
| Old 1 | 86 | Fascia Lata | F |
| Old 2 | 74 | Quadriceps | F |
| Old 3 | 80 | Quadriceps | M |
| Old 4 | 60 | Fascia Lata | M |
| Old 5 | 79 | Quadriceps | M |
| Old 6 | 76 | Quadriceps | F |
| Old 7 | 76 | Quadriceps | F |
| Old 8 | 70 | Quadriceps | F |
| Old 9 | 81 | Tensor Fascia Lata | F |
| Old 10 | 64 | Quadriceps | M |
| Old 11 | 65 | Quadriceps | M |
| Old 12 | 75 | Quadriceps | M |
| Old 13 | 64 | Quadriceps | M |

**a,** Table with all donnors age, muscle type and sex information.
